# Modeling the Relationship between the Capsid Spike Protein Stability and Fitness in ϕX174 Bacteriophage

**DOI:** 10.1101/2025.09.02.673881

**Authors:** Y. Sapozhnikov, J.T. Van Leuven, J.S. Patel, C.R. Miller

## Abstract

**Background:** Protein function depends on the stable folding or binding of peptide chains, and the extent to which an amino acid substitution disrupts this stability can be a strong predictor of the mutation’s impact on the protein’s function and the organism’s fitness. This study seeks to understand this relationship in bacteriophage ϕX174 using an experimental technique of deep mutational scanning and computationally predicted protein stability.

**Results:** Analyzing the viability data using a newly developed multistage binomial model confirms that highly destabilizing mutations predict inviability. For single-site variants, the model predicts that ΔΔG of folding greater than ~3 kcal/mol and ΔΔG of binding greater than ~6 kcal/mol cuts the probability of survival by half of the maximum. The maximum probability, even for minimal ΔΔG values, plateaus at around 87%, reflecting the influence of unobserved factors. Fitting of double-site variants data shows similar results. In contrast, a linear regression analysis of viable mutants reveals that the effect of protein stability is too weak to be a reliable predictor of fitness.

**Conclusion:** In this study, we introduce a novel modeling technique to explore the relationship between protein stability and survival outcomes. While large destabilization often indicates inviability, most destabilizing mutations exert only small to moderate effects and do not reliably predict survival. Although protein stability is a crucial biophysical property influencing biological systems, it represents just one of multiple factors that contribute to fitness and survival.

## Background

The fitness of an organism, from the simplest form such as viruses to the most complex such as humans, hinges on functional proteins carrying out their enzymatic activities and providing structural integrity necessary for life. Proteins must fold into their native form and/or bind to other proteins or ligands to function. Changes in Gibbs free energy (ΔG) is a thermodynamic measure of stability that describes whether or not protein folding/binding is favorable. When ΔG of folding is less than zero, then at equilibrium more protein molecules are in a folded conformation than in unfolded conformation because the folded, native state is thermodynamically more stable. Amino acid substitutions can potentially alter the stability, denoted by ΔΔG, and thus disrupt the protein’s function. While the vast majority of amino acid changes are neutral to slightly destabilizing with respect to stability (1,2), a destabilizing mutation can prevent proper folding or binding. Depending on the nature of the protein’s function, a highly stabilizing mutation can disrupt function as well due to the excessive rigidity.

The mutational effect on protein stability has a wide range of implications in various disciplines. In biomedicine, computational calculation of ΔΔG is harnessed to screen for potentially pathogenic mutations (3,4), as destabilizing mutations are associated with protein misfolding and aggregation resulting in a disease (5–7). On the other hand, conformational flexibility of less stable proteins can potentially facilitate gaining new functions and thus evolutionary adaptation (8–10). The reverse – a novel protein with new functions will tend to be less stable – is also true (2,11), and thus stabilization is critical in the design process in protein engineering field. There is an aspect of epistasis in this stability-function trade-off. A protein may have a mutation that preserves function but destabilizes the protein enough so that a second mutation pushes it beyond the threshold to render it inviable (12). Conversely, a stabilizing mutation may provide the background upon which a greater range of subsequent mutations are tolerated, allowing for a novel evolutionary trajectory (2,13–15).

The scale of studying the mutational effect in the context of protein stability grew tremendously in recent years owing to two developments: 1) the proliferation of computational methods to predict ΔΔG based on protein sequences or structures, 2) the development of deep mutational scanning (DMS) techniques which allow a large-scale assay of protein variants. Laboratory techniques of measuring ΔΔG are typically costly and time-consuming, making it difficult to scale. However, computational methods offer an alternative where large numbers of mutations can be evaluated relatively quickly to obtain predicted ΔΔG values. While many algorithms using different types of force fields exist (16,17), FoldX (18,19) is a popular option due to its speed and ease of use. The performance of these algorithms have been extensively evaluated (4,17,20–23). Also, we previously developed a predictive model to quantify the errors associated with FoldX prediction (24) to compensate for the lack of a measure of uncertainties in the software. On the experimental front, a typical DMS study involves generating a large library of protein variants (i.e., saturation mutagenesis), followed by a functional assay and high-throughput sequencing (25). The technique led to numerous studies of protein variants relevant to human health (26–28), pathogenic viruses (29–32), and meta-analyses of DMS studies, simultaneously analyzing thousands of variants from various protein systems in order to gain generalizable insights (33–35).

In this work, we characterize the stability effect of single and double amino acid substitutions in the major spike protein G of a bacteriophage ϕX174 on the viability of the virus. ϕX174 is a tailless, icosahedral virus with a single-stranded DNA genome and belongs to the *Microviridae* family (36,37). Each monomer of the G protein is 175 amino acids-long and multimerizes into a pentamer that binds with a pentamer of the coat protein F to form the icosahedral capsid (Figure 1A). The G protein is involved in recognition of and attachment to host cells (37). Using a mutagenesis library of the G protein (38), we performed a flask growth experiment and analyzed the viability of 681 single-site mutation variants and 988 double-site mutation variants. We show that there is a critical ΔΔG threshold by which we can predict the viability of mutant phages using a novel statistical model that incorporates the uncertainty of a biological system and experimental data.

**Figure 1.**
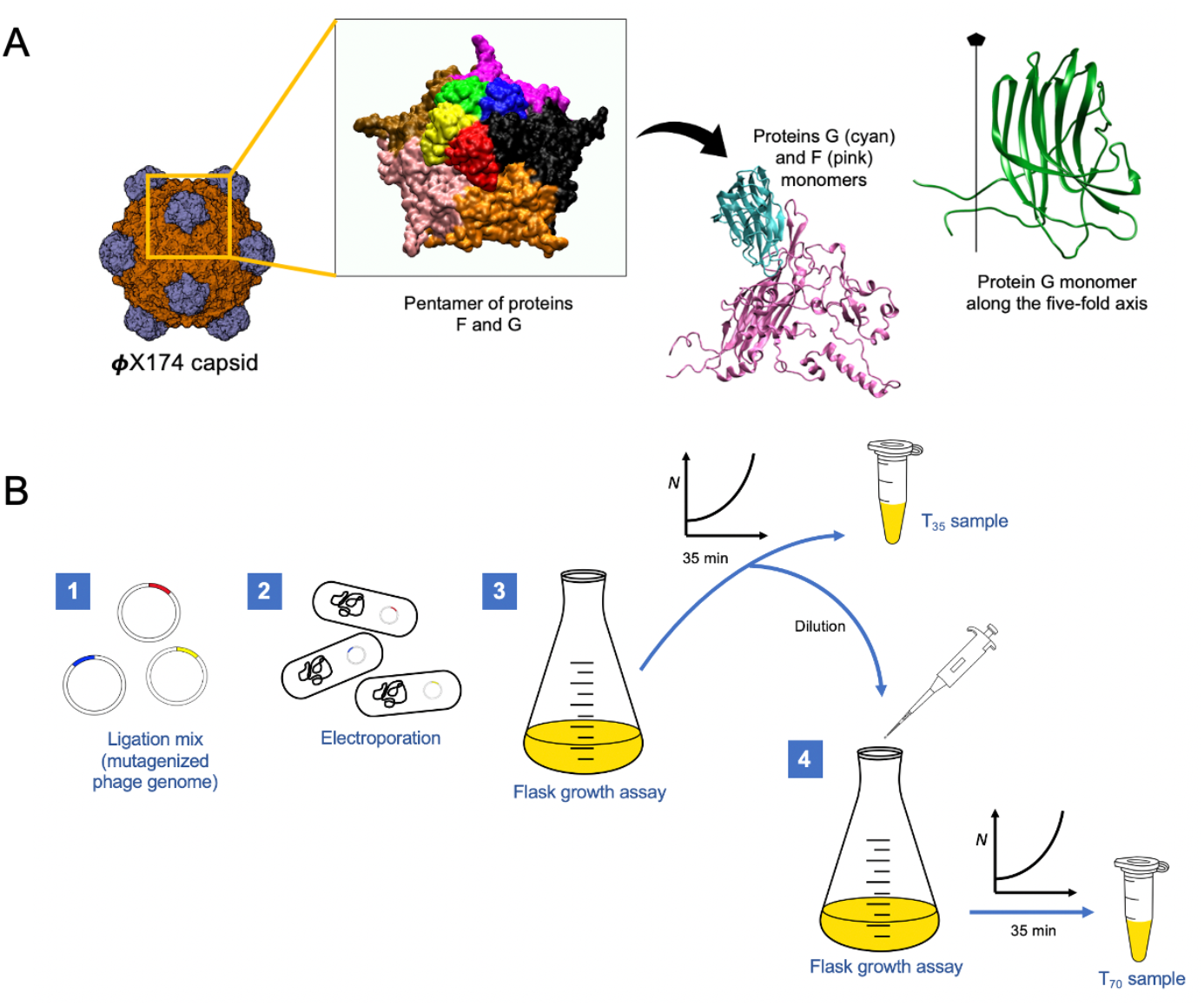
ϕX174 model system and the experimental procedure. (A) The icosahedral capsid consists of 12 pentamers of proteins F and G (60 copies of each protein total). A pentamer of G sits atop a pentamer of F, forming a spike. The image of capsid is provided by the Protein Data Bank: PDB ID 2BPA (49). The F-G pentamer and heterodimer are created using VMD (50). The ribbon representation of a G monomer is adapted from (37) (B) Ligation mix assembled with mutagenic libraries of protein G [step 1] are electroporated into XL1-Blue electroporation-competent cells [step 2]. Phage lysate was added to a flask containing *E. coli* host cells and grown for 35 minutes [step 3] after which a small aliquot is transferred to another flask with host cells and grown for additional 35 minutes [step 4].

## Methods

### Generating mutant virus library

The detailed procedure of generating the ϕX174 mutant libraries is described by Faber and colleagues (38). To summarize, the ϕX174 genome was separated into 14 fragments where each segment is flanked by unique 6 base overhang sequences and cloned into pCR2.1 plasmid vectors. Mutations were introduced into ϕX174 gene F (fragments F1, F2, and F3) and gene G (fragments G1 and G2) via nicking mutagenesis (39). These fragments were assembled via Golden Gate cloning to generate complete circular ϕX174 genomes with mutations in the desired gene fragments. 500ng of each of the 14 plasmids were combined and digested by mixing with 2.5uL of 20mM DTT, 1.5uL of *Bsm*B1, 5uL of 10X gel loading buffer with dye, and water to make 50uL total volume. The mix was incubated at 37°C heat block for 15-30 minutes. All 50uL was loaded into a 1.2% agarose gel to separate DNA by electrophoresis and DNA fragments of length approximately 200-1000 bp were extracted using Thermo Scientific GeneJet Genomic DNA Purification kit. 17uL of the extracted fragments were mixed with 2uL 10X ligase buffer and 1uL T4 DNA ligase (Promega), and incubated in 14°C heat block overnight.

Ligated genomes were electroporated into XL1-Blue electroporation-competent cells according to the manufacturer protocol. Cells were then transferred to culture tubes with 50uL DNase, incubated at 37°C in a shaker for 1 hour. After spinning out the cells, the filtrate was concentrated using Amicon Ultra-0.5 Centrifugal Filter Units, according to the manufacturer protocol and diluted 10uL in 990uL of ϕLB media (10g/L tryptone, 5g/L Bacto yeast extract, 10g/L NaCl, and 2mM CaCl_2_) for use in flask growth assay while the rest was saved for PCR.

Four sets of transformation mix were produced – 1 wild-type as a control and 3 replicates of G protein mutants. Because the ligated genomes were assembled with both G1 and G2 mutagenic libraries, each of which contains single amino acid substitutions, we expected a combination of wild-type, single-site mutants, and double-site mutants in the transformants.

### Flask growth assay

The pool of mutant viruses was then propagated on *E. coli C* (accession number CP024090), which is the typical ϕX174 host used in our lab. Flask growth assay was carried out for each of the transformation mix as follows. Host cells were prepared by growing to ~10^8^ cells/mL, of which 1.5mL was aliquoted into a flask containing 8.5mL of ϕLB in 37 °C water bath. For each set of the transformation mix, an aliquot size was calculated to contain approximately 30,000 phage particles based on titers. This aliquot of phage was added to the flask with actively growing host cells and incubated in the shaking water bath. 200uL of samples were taken from each flask at 35 minutes. An additional 10uL from each flask was transferred to a second flask with actively growing host cells (prepared exactly same as the original assaying flask). After another 35 minutes (70 minutes total), 200uL samples were taken from each flask. All samples were treated with 50uL of chloroform to kill host cells and prepared for PCR using HighPrep PCR cleanup kit (MagBio Genomics).

### Sequencing

We followed Dual Barcoded Two-Step PCR procedure provided by the Genomics Resources Core at the University of Idaho, with some modifications (40). The primers for the first PCR reaction amplify gene G and add ~20 bp for the annealing of barcoding primers (F-ACACTGACGACATGGTTCTACAtctgcttaggagtttaatc, R-TACGGTAGCAGAGACTTGGTCTgcaccaaacataaatcacc). The lowercase sequence anneals to the ϕX174 genome, and the uppercase sequence anneals to the second step primers. A short linker region of variable length (2-5 bp) adds diversity to the library for improved sequencing. The first PCR reaction contained 5uL 10X buffer, 0.8uL dNTPs, 5uL primers, 0.5uL Taq polymerase, 2uL template, and 36.7uL water. Thermocycling conditions were: 1 cycle at 95°C for 1 min, 25 cycles at 95°C for 30 s, 50°C for 30 s, 68°C for 1 min, and 1 cycle at 68°C for 5 min. The second PCR adds the standard P5/P7 Illumina adapter sequences and 6 bp barcodes that enable multiplexing. Each sample and replicate were uniquely barcoded so that sequences could be demuliplexed afterwards. The second PCR reaction contained 5uL 10X buffer, 0.8uL dNTPs, 1.875uL barcoding primers, 0.5uL Taq polymerase, 2.5uL diluted template, and 39.4uL water. The amount of dilution of the first-round products normalizes the templates for the second PCR reactions and is based off of gel electrophoresis. The range in dilution factor is 1:10 to 1:3. Thermocycling conditions were: 1 cycle at 95°C for 1 min, 10 cycles at 95°C for 30 s, 60°C for 30 s, 68°C for 1 min, and 1 cycle at 68°C for 5 min. Then, 2uL of each PCR product was pooled together. Sequencing on Illumina MiSeq and demultiplexing were performed by the IIDS Genomics and Bioinformatics Resources Core at the University of Idaho following the dbcAmplicons protocol (github.com/msettles/dbcAmplicons).

### Sequence data processing and scoring

Sequences underwent quality check and trimming using default parameters in fastp (41) and paired end reads were merged using FLASH (42). We used the denoising algorithm in DADA2 (43) to correct sequencing errors and remove singletons. Mutation frequencies were counted using an ad-hoc R script that mapped reads using BWA (44) then enumerated mutations by parsing CIGAR strings from mapped reads. Enrich2 (45) was used to calculate fitness scores.

### Obtaining predicted ΔΔG of folding and binding stability

Crystal structure of ϕX174 capsid was downloaded from RCSB Protein Data Bank (PDB ID: 2BPA) and modified to remove everything except pentamers of G and F proteins. Missing atoms were added using WHAT IF web server (https://swift.cmbi.umcn.nl). Structures of a G monomer and a trimer were prepared and subjected to MD simulations using GROMACS 5.0.7. Details of our protocol for MD simulation and FoldX are described in our previous studies (46,47). Briefly, simulations were run for 100ns during which 1ns snapshots were taken. For each snapshot, we used RepairPDB command in FoldX to minimize potential energy and obtain its convergence, followed by BuildModel command to generate all possible 19 single amino acid mutations at each residue. Mutational effects on folding stabilities (ΔΔG_fold_) were determined using Stability command on the monomer structures while AnalyseComplex command was applied to the trimer complexes for effects on binding stabilities (ΔΔG_bind_). We then calculated final ΔΔG_fold_ and ΔΔG_bind_ for each mutation by averaging the FoldX estimates across all 100 individual snapshot estimates. For double-site variants, we assumed an additive effect and summed the ΔΔG_fold_ or ΔΔG_bind_ values of constituent mutations. Any doubles containing nonsense or synonymous mutations were removed for further analyses.

### Statistical analysis

All statistical analyses were performed in R (version 4.2). In order to model the relationship between ΔΔG and viability, we used our novel Multistage Binomial Model (MBM) (48), an extension of the standard logistic model. MBM models phenomena that occur as a series of stages where a failure in one stage results in a failure as a whole, i.e., multiplicative risk. This conceptualization represents many biological processes more realistically than with traditional logistic model. For example, a peptide chain must fold properly first, and then can bind other subunits or ligands. Failure in either folding or binding will knock out the protein function. MBM can also accommodate effects of unobserved factors on the binary outcome, by allowing a maximum or minimum probability that may be less than 1 or more than 0, respectively. By contrast, the traditional logistic model makes the often invalid assumption that the success probability goes to 1 or 0 as the predictor variable(s) in the model change. If the available data indicates that the specified variables are not enough to determine the outcome definitively, the success probability in MBM will approach an asymptote that is less than 1 or greater than 0. Lastly, MBM can incorporate measurement errors. This means that datapoints with large uncertainties will carry less weight than datapoints with smaller uncertainties. We have previously developed a predictive model to estimate potential errors of FoldX output (24). We obtained the uncertainties of ΔΔG_fold_ and ΔΔG_bind_ for our variants using the error model, and incorporated them into MBM.

We categorized variants that were observed only at time zero (*T*_0_) as “ inviable,” and those that were observed at later time points regardless of their presence at *T*_0_ as “ viable.” The data were then fitted to MBM with this viability category as the binary response variable, and ΔΔG_fold_ and ΔΔG_bind_ as covariates. We further analyzed the viable variants using multivariate linear regression model.

## Results

### Flask growth assay result

The time zero (*T*_0_) sample and the three replicates of 35-minute (*T*_35_) and 70-minute (*T*_70_) samples yielded a total of 757,837 reads, and 2028 unique variants and wild-type. Among these, single-site variants consisted of 382,819 reads and 688 unique variants (Table 1). The number of unique amino acid substitutions at each residue varied across the span of protein G (Figure 2B).

**Table 1.**
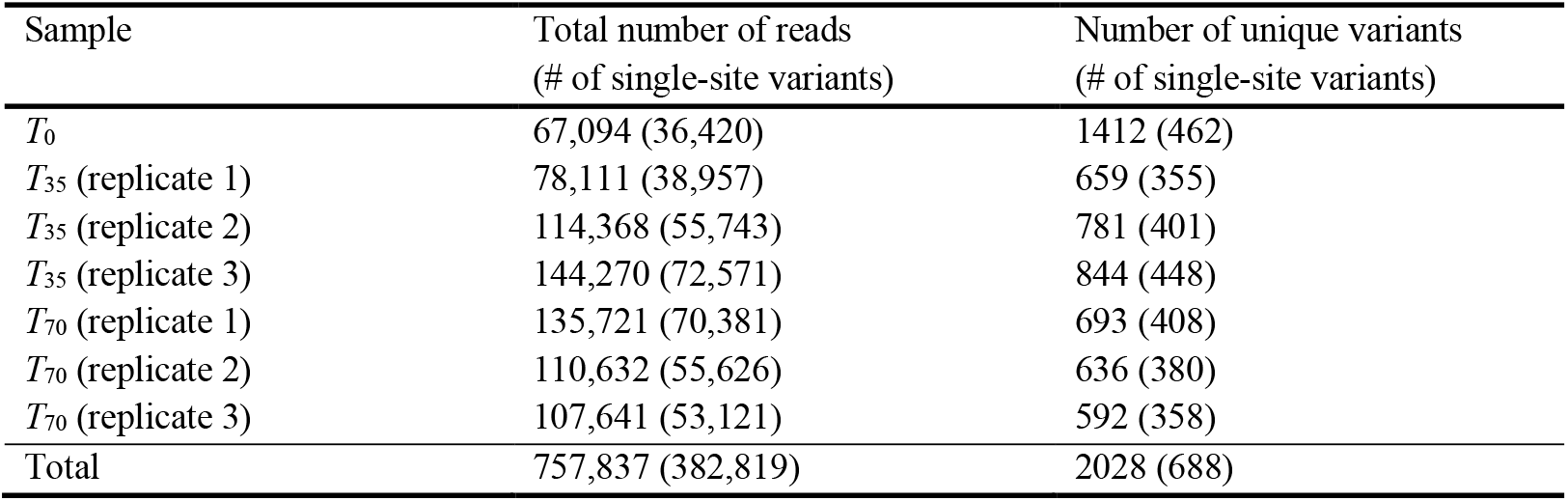
Sequencing of phage variants. Pooled samples yielded a total of 757,837 reads comprising of 2028 unique variants and wild-type. Demultiplexed counts for individual samples are shown.

| Sample | Total number of reads<br>(# of single-site variants) | Number of unique variants<br>(# of single-site variants) |
| --- | --- | --- |
| $T_0$ | 67,094 (36,420) | 1412 (462) |
| $T_{35}$ (replicate 1) | 78,111 (38,957) | 659 (355) |
| $T_{35}$ (replicate 2) | 114,368 (55,743) | 781 (401) |
| $T_{35}$ (replicate 3) | 144,270 (72,571) | 844 (448) |
| $T_{70}$ (replicate 1) | 135,721 (70,381) | 693 (408) |
| $T_{70}$ (replicate 2) | 110,632 (55,626) | 636 (380) |
| $T_{70}$ (replicate 3) | 107,641 (53,121) | 592 (358) |
| Total | 757,837 (382,819) | 2028 (688) |

**Figure 2.**
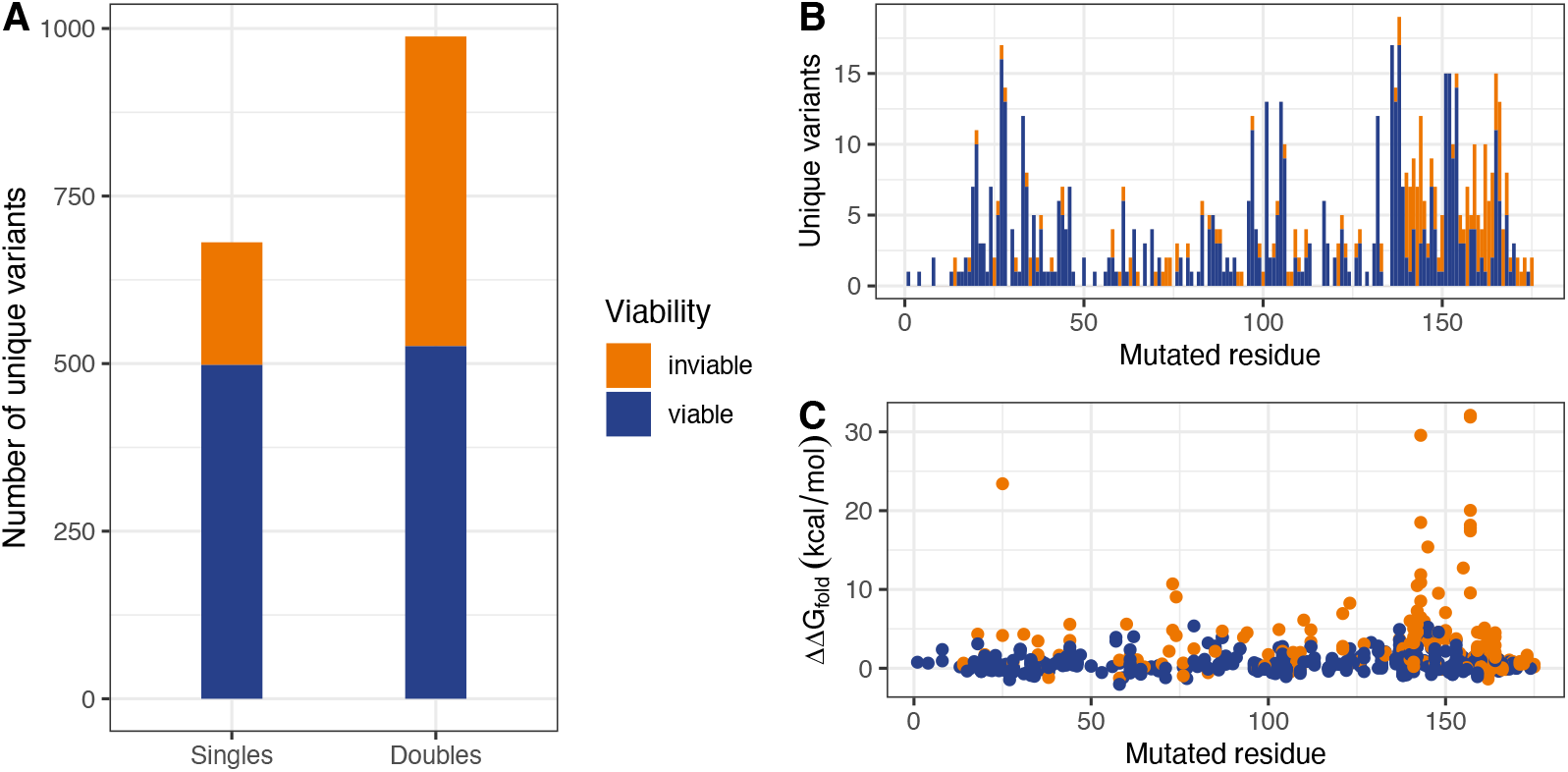
Characterization of observed variants. (A) Number of observed variants containing one or two mutations from wild-type. Variants that were observed in the *T*_0_ sample, but no subsequent samples were considered inviable. Among 681 observed single-site variants, 498 (73%) were viable and 183 (27%) were not. Among 988 double-site variants, 526 (53%) were viable and 462 (47%) were not. (B) The histogram shows varying distributions of unique variants per residue. (C) The scatterplot shows viable variants tend to have neutral or moderately destabilizing mutations (small ΔΔG_fold_) while many inviable variants have highly destabilizing mutations (large ΔΔG_fold_).

### Highly destabilizing mutations predict inviability

Of the 688 variants with single amino acid substitutions that were analyzed, 184 (27%) variants were only observed at *T*_0_ and thus classified as inviable. 504 variants (73%) were observed in subsequent time points at least once, irrespective of their presence at *T*_0_, and thus classified as viable. A scatterplot of single-site variants along their residue position and ΔΔG_fold_ shows that viable variants tend to have low ΔΔG (i.e., predicted to be neutral to moderately destabilizing). In contrast, variants with high ΔΔG (i.e., predicted to be highly destabilizing), tend to result in inviable particles even if they do transform.

In our previous work with molecular modeling of protein G, substitutions of Gly119 have shown to be problematic due to the extremely large Van der Waals clash that results from mutating glycine residues. The errors in FoldX predictions increase with large Van der Waals clash (24), and the inaccurate prediction is particularly pronounced at residue 119. Therefore, we excluded 7 variants involving Gly119 from subsequent stability analyses.

Analysis of viability with ΔΔG shows that variants with high ΔΔG are less likely to be viable. MBM output (Table 2) shows that each unit increase in ΔΔG_fold_ results in 1.4 *decrease* in log odds of survival while each unit increase in ΔΔG_bind_ results in 0.4 *decrease* in log odds of survival. At approximately 3 kcal/mol of ΔΔG_fold_ or 6 kcal/mol of ΔΔG_bind_, we can expect less than half of the maximum probability of survival (Figure 3). The maximum probability of survival as predicted by MBM is 0.87 (Table 2). This means that no matter how small ΔΔG, we do not predict a 100% survival.

**Table 2.** MBM fitting result. MBM was fit to the viability and ΔΔG data with measurement errors predicted by the FoldX error model (24). For double-site variants, ΔΔG values for each constituent mutation were summed under additive assumption. Estimates of intercepts (ϕ_0_) and coefficients (ϕ_1_) for each predictor variables and the range of the probability of being viable (success probability limits) are shown.

| Parameter |  |  | Estimate | p-value |
| --- | --- | --- | --- | --- |
| Single-site variants | $\Delta\Delta G_{\text{fold}}$ | $\beta_0$ | 5.2172 | <0.001 |
| | | $\beta_1$ | -1.4047 | <0.001 |
| | $\Delta\Delta G_{\text{bind}}$ | $\beta_0$ | 2.7108 | <0.001 |
| | | $\beta_1$ | -0.3850 | <0.001 |
|  | Success probability limits |  | (0, 0.8676) | - |
| Double-site variants | $\Delta\Delta G_{\text{fold}}$ | $\beta_0$ | 8.1501 | <0.001 |
| | | $\beta_1$ | -1.9123 | <0.001 |
| | $\Delta\Delta G_{\text{bind}}$ | $\beta_0$ | 4.8494 | <0.001 |
| | | $\beta_1$ | -1.1085 | <0.001 |
|  | Success probability limits |  | (0, 0.7851) | - |

**Figure 3.**
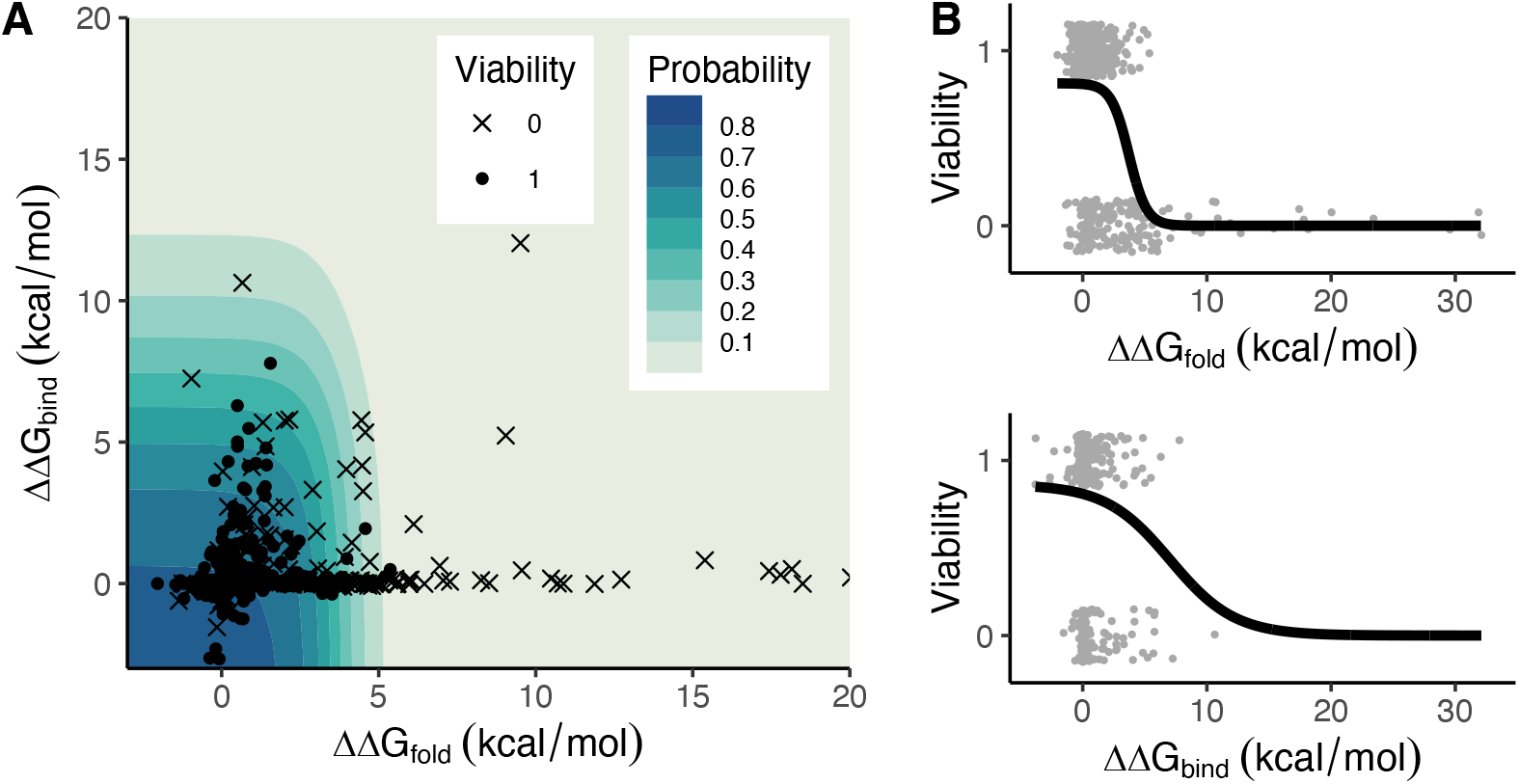
Viability of single-site variants predicted by multistage binomial model. (A) ΔΔG of folding and binding of 681 variants are plotted over the contour plot indicating the probability of survival. Solid dots indicate viable mutants while crosses indicate inviable mutants. Data points beyond 20 kcal/mol are not shown. (B) The probability of being viable as ΔΔG of folding (top) or binding (bottom) increases while the other variable is held constant. The probability never reaches one, reflecting that there are other unobserved biological processes in addition to ΔΔG affecting the outcome.

### Neutral or moderately destabilizing mutations do not correlate with fitness

Fitting linear regression models to viable variants’ data (n = 498) revealed a weak, negative relationship between ΔΔG and fitness score: the coefficients for ΔΔG_fold_ and ΔΔG_bind_ in univariate models are −0.15 and −0.09, respectively (Figure 4), which remain similar in the multivariate model (Table 3). *R*^*2*^ is slightly greater in the multivariate model than univariate models. Altogether, the small effects and negligible *R*^*2*^ values indicate that ΔΔG is not an effective predictor for viral fitness.

**Table 3.**
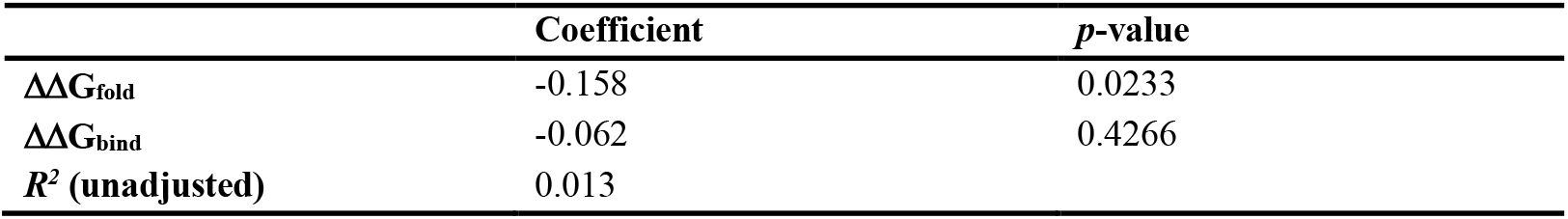
Multiple linear regression model details. (A) Multiple linear regression of fitness with ΔΔG_fold_ and ΔΔG_bind_ as predictor variables show negative effects of ΔΔG on fitness. However, the effect of ΔΔG_bind_ is not statistically significant and overall *R*^*2*^ is small.

**Figure 4.**
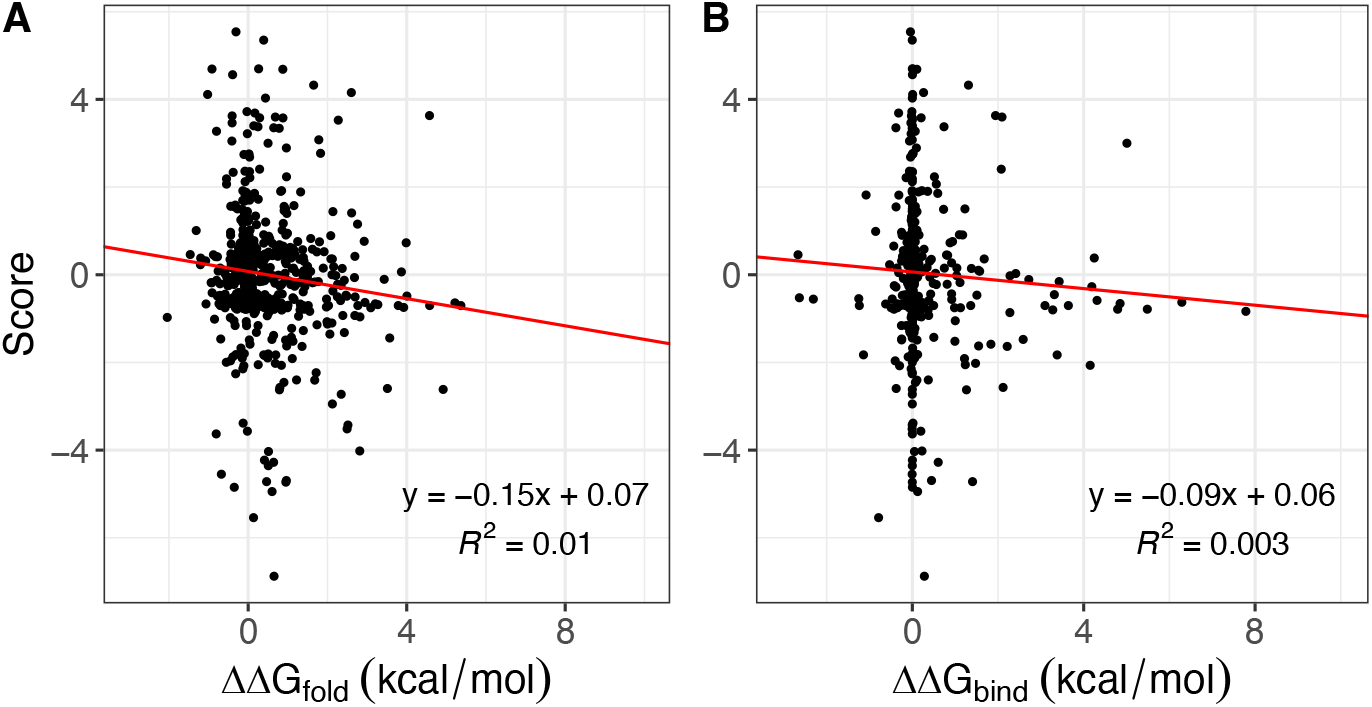
Univariate linear regression between ΔΔG and fitness scores in single-site variants. (A) ΔΔG_fold_ of most variants are clustered near 0 kcal/mol with a wide range of fitness scores. While ΔΔG_fold_ has an overall negative effect on fitness, the association is weak. (B) A scatterplot of ΔΔG_bind_ and fitness scores also show a weak, negative relationship. Clustering near 0 kcal/mol is more pronounced with ΔΔG_bind_.

### Viability of double-site variants can be predicted by combined stability

For variants with two mutations, there were 988 such variants after filtering (see Methods). Among them, 285 variants contained mutations that were not observed in the single-site variant data while 703 variants contained one or both mutations that were found in the single-site variant data and thus their viability is known. Two double-site variants in which both constituent singles were known to be inviable turned out to be inviable, while 214 out of 236 double-site variants that had both constituent singles known to be viable turned out to be viable (Table 4, Figure 5B). In cases where the viability of only one of the two mutations is known, the vast majority (66 out of 72) of the double-site variants that have at least one inviable mutation were inviable (Table 4).

**Table 4.**
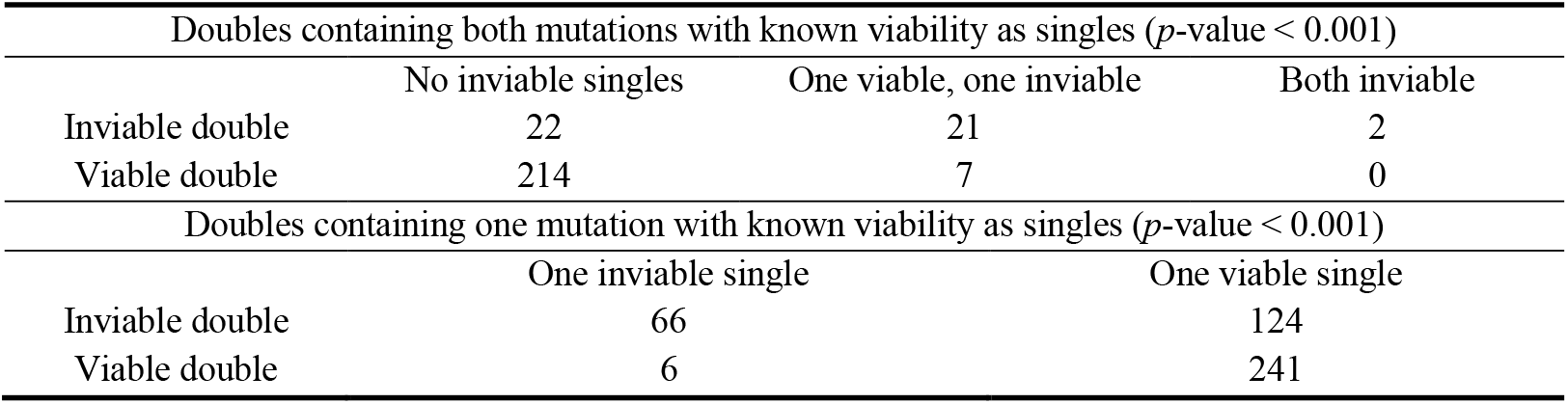
Counts of double-site variants with viability of constituent mutations. There were 266 double-site variants containing both mutations with known viability because they were observed as single-site variants. 437 variants contained one of the two mutations with known viability. The counts are broken down according to the doubles’ viability and the constituent mutations’ viability. *p*-values were calculated by chi-square test for independence.

**Figure 5.**
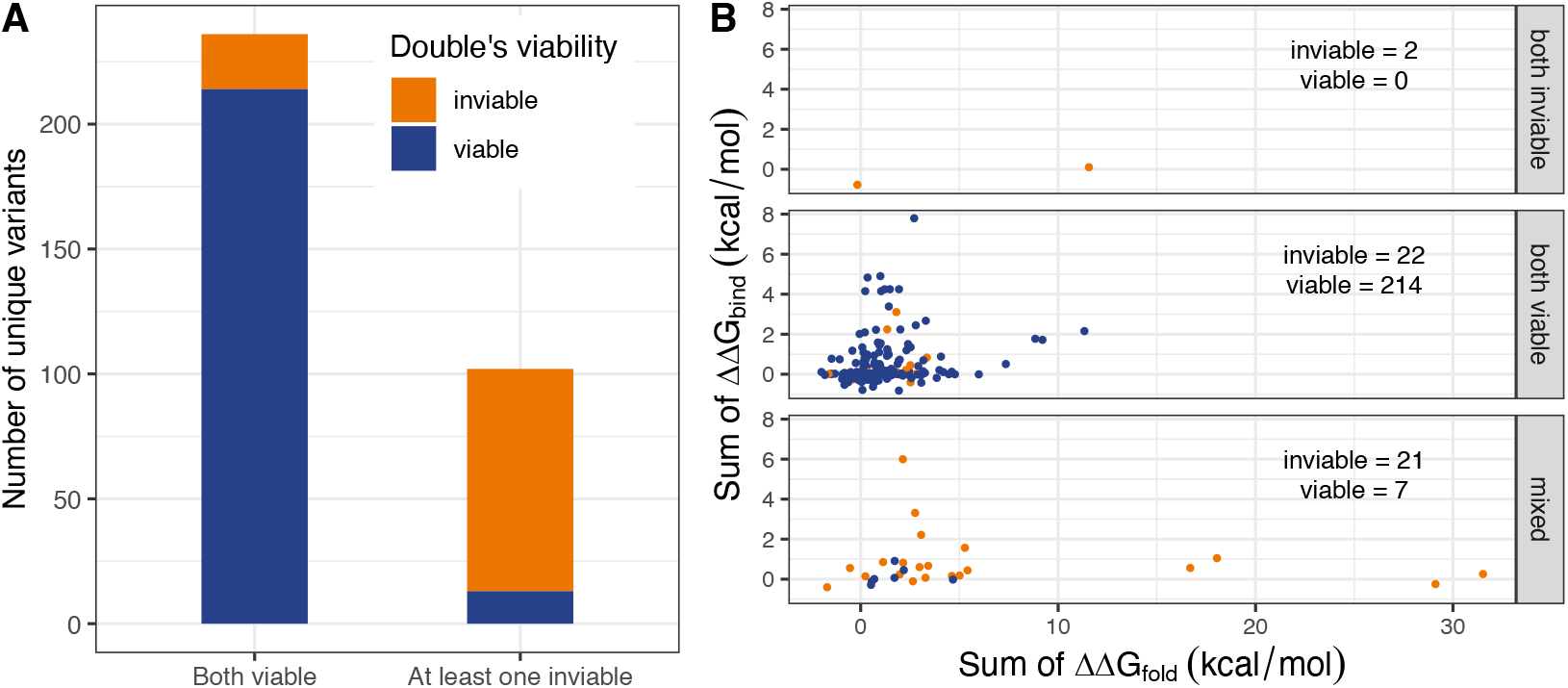
Viability of double-site variants. (A) Among 236 double-site variants that had both viable mutations, 214 (91%) were viable and 22 (9%) were inviable. Among 102 double-site variants with at least one inviable mutation, 13 (13%) were viable and 89 (87%) were inviable. (B) 266 double-site variants had both mutations of known viability. Their viability as a double and summed ΔΔG values are plotted, separated in each panel according to the viability of constituent mutations.

As with the single-mutation viability data, we fit the double-mutation data to MBM with combined ΔΔG_fold_ and ΔΔG_bind_ of constituent mutations as predictors. The model shows that each unit increase in combined ΔΔG_fold_ results in 1.9 *decrease* in log odds of survival and each unit increase in combined ΔΔG_bind_ results in 1.1 *decrease* in log odds of survival (Table 2). For both ΔΔG_fold_ and ΔΔG_bind_, the probability of survival becomes less than half of the maximum probability beyond approximately 4 kcal/mol (Figure 6).

**Figure 6.**
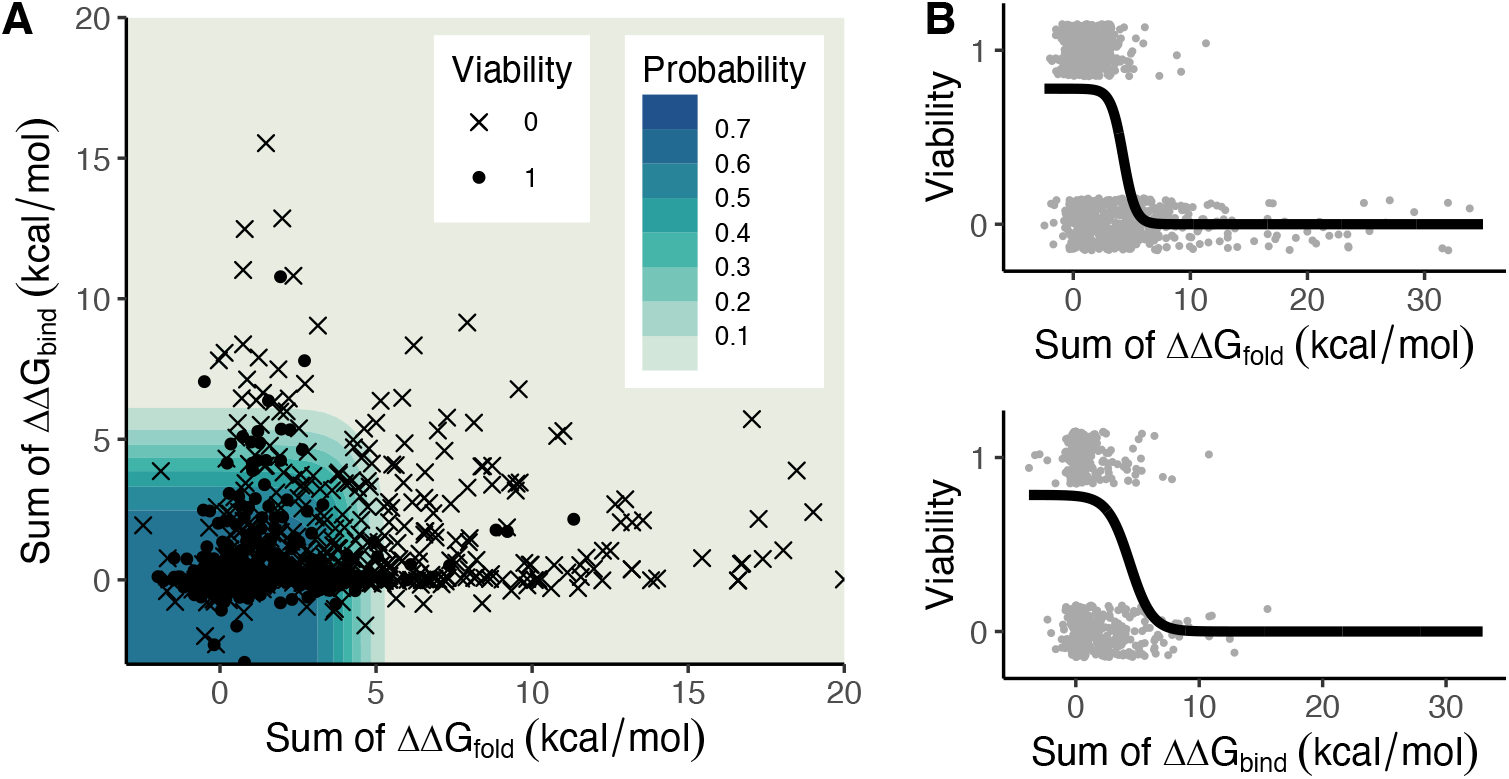
Viability of double-site variants predicted by multistage binomial model. (A) 988 double-site variants are plotted with ΔΔG_fold_ and ΔΔG_bind_ of their constituent mutations combined. The contour plot indicates the probability of survival as predicted by MBM. Solid dots indicate viable mutants while x indicates inviable mutants. Data points beyond 20 kcal/mol are not shown. (B) The probability of being viable as combined ΔΔG_fold_ (top) or ΔΔG_bind_ (bottom) increases while the other variable is held constant.

## Discussion

A DMS approach produces large-scale mutational data that can elucidate functional consequences of every possible amino acid substitutions at each residue (25). We apply this approach to bacteriophage ϕX174 to investigate the mutational effect on the stability of a capsid protein and the consequences on the fitness and survival of the phage. This is the first time that DMS approach was used on ϕX174 to our knowledge.

In order to systematically describe the relationship between protein stability and mutant phages’ viability, we developed MBM (see Methods), a modified logistic regression method that models multistage processes inherent in biological phenomena. The modeling result provides an insight into the stability threshold that can be tolerated. In single-site variants data, we see that the probability of survival drops below half of the maximum probability at around ΔΔG_fold_ of 3 kcal/mol or ΔΔG_bind_ of 6 kcal/mol. Once ΔΔG_fold_ is beyond 5 kcal/mol or ΔΔG_bind_ is beyond 10 kcal/mol, the probability is almost zero (Figure 3). The model output and probability curves are similar for double-site variants as well (Table 2, Figure 6), implying that there is a maximum level of destabilization that can be tolerated whether from a single mutation or additive effects from multiple mutations. For ΔΔG values below this threshold, the variants’ fate is more variable. In doubles, the distribution of combined ΔΔG values for the majority of them containing mutations of known viability are similar between viable and inviable doubles (Figure 5B). Regardless of ΔΔG values, simply containing at least one inviable mutation renders the double-site variants more likely to be inviable compared to those with both viable mutations (87% vs 9%, Figure 5A).

An important feature of MBM is that the model allows the maximum (or minimum) success probability to approach an asymptote other than 1 (or 0). In a standard logistic regression, the link function transforms predictor values into probabilities between 0 and 1. The model assumes that we can expect a certainty of a binary outcome given a sufficiently large or a sufficiently small predictor value. This assumption is problematic in our viability data. While we can expect a variant to be inviable given a very large ΔΔG_fold_ or ΔΔG_bind_, the reverse is not true as noted above; most variants’ ΔΔG values cluster near 0 kcal/mol and the viability outcome of those variants can be in either direction (Figures 3 and 6). A vast majority of missense mutations are neutral or destabilizing (1,2,25) and ΔΔG of the destabilizing mutations tend to be small. In our dataset, a small number of mutations with large ΔΔG seems to drive the negative trend of decreasing fitness. However, this effect is weak because of the variants with neutral to weakly destabilizing mutations span a wide range of fitness. Therefore, the mutational effect on protein stability does not adequately explain the mutational effect on fitness, which is likely from unknown underlying biological processes. Mutations in protein G may disrupt interactions with coat protein F (36) or external scaffolding protein D (51) without increasing ΔΔG_fold_ or ΔΔG_bind_ in our consideration. Mutations may also affect how the phage recognizes host cells. The influence of these possible biological factors on viability are captured in MBM as the maximum success probability that does not reach 1. In other words, we mathematically demonstrate that while large ΔΔG is sufficient to predict inviability, small ΔΔG is necessary but not sufficient to predict viability due to unobserved factors.

Recently, a growing number of published DMS datasets enabled large-scale analyses investigating generalizable characteristics of amino acid substitutions. A 2022 study by Høie and colleagues, in particular, demonstrated a global relationship among fitness, evolutionary conservation, and thermodynamic stability using a large dataset of more than 150,000 variants collected from 29 protein systems (34). As we have seen in our ϕX174 data, the study showed that variants with low ΔΔG had both high and low fitness, indicating that small perturbation of stability was not a reliable predictor of fitness. The authors also analyzed the fitness against multiple-sequence-alignment-based conservation scores and showed that the conservation score was strongly correlated with fitness (34). It is not surprising that a measure of conservation performs very well as a predictor of fitness since it encapsulates all known and unknown biological factors that led to the evolutionary success, and therefore has been used in various tools for predicting the effect of mutations (52,53). Extant genomes are products of nature’s exploration of sequence space by way of evolution, but we do not know the exact paths of substitutions that have occurred and why many of them fail. As fundamental as it is at a biophysical level, protein stability is only one of multiple factors that determine viability and fitness.

The limitation of our study includes stochasticity inherent in the experimental design and in sampling. First, not all mutagenized and ligated genomes enter the cells during the electroporation. Second, a sampling error can lead to an incomplete sequence data; variants with very low fitness may not grow in sufficient numbers and thus fail to be sampled, resulting in incorrect labeling as inviable. However, the strength of our design is the large of number of variants being studied. While the DMS study design may lack the precision at an individual variant level, the large number of variants as a whole strengthens the statistical inference.

## Conclusions

We combined an experimental technique of DMS and computational method of predicting stability changes to study the relationship between capsid protein stability and fitness in ϕX174 system. Using a newly developed multistage binomial model, we showed predicted ΔΔG greater than 5 kcal/mol in single-site variants and double-site variants of protein G tends to render the phages inviable. However, ΔΔG was not a statistically significant predictor of fitness in viable mutants. From our results as well as other DMS studies, we can conclude that highly destabilizing mutation is detrimental to fitness while stabilizing, neutral, or moderately destabilizing mutations do not show definitive correlations with fitness – likely due to numerous unobserved biological processes that affect fitness.

## Notes

### Competing Interest Statement

The authors have declared no competing interest.

